# A resource and computational approach for quantifying gene editing allelism at single-cell resolution

**DOI:** 10.1101/2025.05.22.653824

**Authors:** Matthew H. Ung, Gabriella Angelini, Ruijia Wang, Alyssa Pyclik, John R. Lydeard, Juliana Xavier Ferrucio, Michelle I. Lin, Tirtha Chakraborty, Huanying Gary Ge

**Affiliations:** Vor Bio, 100 Cambridgepark Dr #101, Cambridge, MA 02140

**Keywords:** Single-cell DNA sequencing, Allelism, Mixture models, Gene editing, CRISPR-Cas9

## Abstract

CRISPR-Cas9-based gene editing is a powerful approach to developing gene and cell therapies for several diseases. Engineering cell therapies requires accurate assessment of gene editing allelism because editing patterns can vary across cells leading to genotypic heterogeneity. This can hinder development of robust cell therapies. Droplet-based targeted single-cell DNA sequencing (scDNAseq) has been used to genotype targeted loci across thousands of cells enabling high-throughput assessment of gene editing efficiency. Here, we constructed a “ground truth” gene editing single-cell DNAseq atlas, along with an artifact-aware computational workflow called GUMM (Genotyping Using Mixture Models) to systematically infer single-cell allelism from these data. This resource was created by expanding CRISPR-Cas9-edited HL-60 clones that harbored distinct insertion-deletion (indel) profiles in *CLEC12A* and mixing them at pre-defined ratios to create artificial cocktails that mimic the potential editing diversity of a CRISPR-Cas9 experiment. This enabled assessment of technical artifacts that confound interpretation of allelism in the readouts of gene edited cells. GUMM was able to accurately genotype cells and infer the original clonal composition of the artificial cocktails even in the presence of artifacts.

## Introduction

The advent of genome editing technologies, notably CRISPR-Cas9-based systems, has ushered in a transformative era in molecular biology. These powerful tools enable precise modification of genes, offering unprecedented opportunities to decipher gene function, study disease mechanisms, and develop novel therapeutic strategies. As gene editing techniques become widespread across various research fields and clinical applications, the imperative of comprehensively assessing allelism in gene-edited cells becomes increasingly evident^1^.

Identifying allelism, the characterization of allelic variants, is fundamental to understanding the consequences of gene editing especially when complex genotypes are desired. Accurate allelic profiling not only confirms the success of intended genetic alterations but also aids in elucidating potential off-target effects^2^ and unintended phenotypic consequences. This knowledge is pivotal for ensuring the safety and efficacy of gene-editing approaches, particularly when applied in the context of gene therapy and regenerative medicine. The primary approach to allelism assessment involves isolating and expanding clones from the edited sample. Clones are then individually selected and profiled with DNA sequencing or PCR to validate the presence of the intended edit. Despite this being a widely adopted standard procedure, it remains time-consuming, low throughput^3,4^, and may not result in an isogenic readout^5^. Innovative solutions have emerged to overcome this bottleneck, such as microfluidic-based targeted single-cell DNA sequencing (scDNAseq). scDNAseq has the capability to barcode and sequence tens of thousands of individual cells within a sample in a relatively brief period, offering a more comprehensive depiction of editing outcomes^6^. Given its ability to provide a readout at single-cell resolution, this technique is ideally suited for profiling cells with intricate genotypes across multiple loci, thereby unveiling heterogeneity of editing consequences.

Due to its high throughput capabilities, scDNAseq requires sophisticated data analysis strategies to decouple signal from noise. Moreover, these strategies must align with experimental objectives and the specific conclusions we aim to derive from the data. In this study, we performed a deep characterization of data derived from scDNAseq of CRISPR-Cas9 edited samples and developed a computational workflow tailored to evaluating allelism in the context of gene editing. Specifically, we created a ‘ground truth’ data atlas by running scDNAseq on artificial cocktails formed by mixing edited HL-60 clones with pre-defined edited allele variants of *CLEC12A*, two pivotal markers within the hematopoietic myeloid lineage. This data resource was used to delineate technical artifacts that could confound downstream interpretation of editing allelism.

Lastly, we leveraged this resource to develop a novel computational workflow called GUMM (Genotyping Using Mixture Models) that systematically genotypes single-cells from scDNAseq data by fitting a series of Gaussian mixture models (GMMs) to allele read counts generated by CRISPResso2^7^. It is uniquely well-suited for analyzing gene editing experiments where cells in the sample are primarily isogenic and differ only by their edited allele profiles. GUMM outputs a probabilistic prediction of cell genotype while being aware of technical artifacts including low coverage at the editing site, PCR amplification imbalance^8^, multiplets (also known as doublets)^9^, and sequencing error^10^. This is distinct from previous methods that specialize in genotyping and identifying multiplets in cell populations with distinct genetic backgrounds^11^. When applied to our ground truth dataset, GUMM was able to rapidly evaluate allelism in single-cells and accurately estimate the original clonal ratios of the artificial cocktails. Our study provides both a novel bioinformatic solution and rich data resource for researchers in the gene editing community looking to engineer cells with complex genotypes.

## Materials and Methods

### CRISPR-Cas9 editing and expansion of HL60 monoclones

HL-60 cells (CCL-240™, ATCC) were cultured in 20% FBS in Iscove’s Modified Dulbecco’s Medium (IMDM, Cat. No. 12440053, ThermoFisher Scientific). Cas9-RNPs were delivered via electroporation using the Lonza Amaxa 4D-Nucleofector System (Cat No. AAF-1002, Lonza Bioscience) to 1e6 HL-60 cells in 100 uL 4D-Nucleofector Sinlge Cuvettes (Cat. No. AXP-1003, Lonza Bioscience) with the SF Cell Line 4D-Nucleofector X Kit L (Cat. No. V4XC-2012, Lonza Bioscience). Post-electroporation, cells recovered in culture for 48 hours. Edited cells were single-cell dispensed into one well of a flat bottom, tissue culture treated 96-well plate with 100 uL of 20% FBS in IMDM using the Namocell Hana Single-cell Dispenser (Cat. No. NI004, Namocell). Cells were expanded to confluency and genotyped using Sanger sequencing followed by ICE Analysis.

### Generation and validation of cocktails

Monoclonal singleplex edited HL-60 cell lines were mixed at defined proportions with one another and unedited HL-60 cells that were cultured for the same amount of time as the monoclonal edited cells. To generate the cocktails, each cell line was counted in duplicate using the Nexcelom Cellometer (Auto 2000, Nexcelom) and the average total number of cells was used to calculate the number of cells to add to the mixture. Cocktails were generated immediately before running the MissionBio Tapestri protocol.

### scDNAseq of artificial cocktails

We produced barcoded single-cell libraries for each cocktail using Mission Bio’s Tapestri platform and a panel of 21 amplicons including two that covered *CLEC12A* editing site. Sample preparation was performed using Mission Bio’s recommended protocol. Cells from the 35% Hom, 55% Het, 10% WT sample were filtered with a 40 uM Flomi before cell encapsulation to generate data with a low multiplet rate. Cells from the 55% Hom, 35% Het, 10% WT and 45% Hom, 45% Het, 10% WT samples were not filtered to generate data with a high multiplet rate. Cocktail libraries were sequenced on Illumina’s NextSeq 2000 with a P2 600 cycle kit.

### Analysis of scDNAseq gene editing data resource

Raw fastq files from each cocktail were processed using the command line implementation of the Tapestri Pipeline (v2.0.2) which performs QC, read trimming, alignment to the reference genome (GRCh38.p14), and barcode deconvolution. The summary report produced by the pipeline was used to assess amplicon uniformity, proper read alignment, and coverage. The pipeline outputs BAM files with each read assigned to a cell barcode under the read group (RG) tag. All BAM files were manually inspected on the genome browser (Qiagen OmicSoft Studio V11.2) to confirm the presence of expected editing patterns at the pseudobulk and single-cell level. To quantify editing, we split each BAM file into individual cell-level BAM files with bamtools. CRISPResso2 was run on each individual cell in “WGS” mode with the following parameters: -- quantification_window_center -3 --quantification_window_size 10 -- min_reads_to_use_region 5 --demultiplex_only_at_amplicons --ignore_substitutions -- exclude_bp_from_left 1 --exclude_bp_from_right 1. The target amplicon regions are slim to the regions of spacer guide RNA with either ±30 bp flanking regions for the internal and public datasets to reduce the effect of variant read length and increase computational efficiency. The output for each cocktail was concatenated into a single table summarizing allele read counts for each barcode. Detailed information on each allele is provided including editing status, DNA sequence, and indel profile. Barcodes with <10 total counts were removed from the data prior to genotyping and sample composition estimation. Prior to GUMM analysis, allele data were collapsed by ignoring substitutions and summing up their counts. Allele frequencies for each cell were calculated by dividing the number of counts from an allele by the total number of counts across all alleles per locus per cell.

### Gaussian mixture modeling of droplet allele frequencies and sample composition estimation

GUMM workflow involves fitting a series of three GMMs to allele frequency distributions in a stepwise manner to genotype individual cells and flag multiplets. GMMs are a class of unsupervised machine learning algorithms widely used to classify data by fitting a finite set of Gaussian distribution models to data assumed to contain distinct clusters of observations. In our application, we apply GMMs to classify droplets based on transformations of their allele counts. Standard GMMs can be described by the following probability density functions:

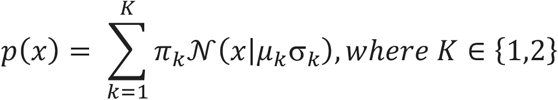

for the univariate case and

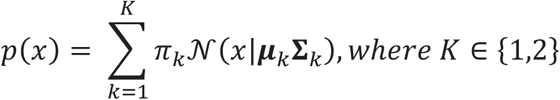

for the multivariate case. K is the number of components or clusters in the data and is restricted to 1 or 2 in our implementation. *π* is the mixing proportion specifying the relative proportions of droplets in each cluster.

GUMM begins by fitting a skewed univariate GMM to the dominant allele frequencies calculated for all droplets using one or two mixing components (i.e. k=1, 2). Skewed GMMs include an extra parameter that describes the amount of skew in the data. A detailed overview can be found in Prates et al. In practice, we found that these skewed models are more accurate when identifying homozygous droplets because dominant allele frequencies of droplets are biased towards 100%. Hartigan’s dip test was used on the ploidy scores to determine *K*where:

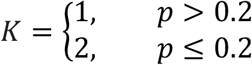

A K=1 model indicates the sample is homogenous (e.g. 100% edited or WT) and a K=2 model indicates a sample is heterogeneous. If K=2, the cluster corresponding to homozygous cells was identified by:

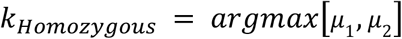

The remaining cells are then used as input for the second step where GUMM flags transparent multiplets by analyzing the ploidy of these non-homozygous droplets at the target locus. Ploidy is assessed by taking the log_2_ allele count ratio of the 2^nd^ and 3^rd^ most common alleles in each droplet. We refer to this ratio as the ploidy score:

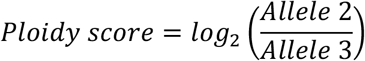

Droplets with only two detectable alleles were whitelisted as true heterozygous cells and were not modeled. A standard univariate GMM was then fit to the ploidy scores to classify the remaining droplets where *x* is the ploidy score of the *i*_th_ droplet and *μ*_k_ and *σ*_*k*_ are the mean and standard deviation of the ploidy score, respectively, of the *k*th component. Again, Hartigan’s dip test was used to determine K because multiplets cannot be detected in pure samples nor do they affect the sample composition estimate. If K=2, the cluster with the smallest mean ploidy score was labeled as transparent multiplets:

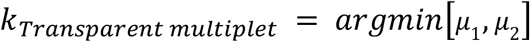

In the third and final step, GUMM classifies the final pool of droplets as either true heterozygous cells or opaque multiplets. This procedure assumes that two alleles that strongly co-occur comprise the genotype of true heterozygous cells. Likewise, droplets with rare allele combinations are most likely erroneous diploid multiplets consisting of two homozygous cells encapsulated in the same droplet (i.e. WT and +1Ins). To leverage this information, GUMM sums up read counts for all alleles across droplets and identifies rare alleles that account for <1% of all read counts. Rare alleles are collapsed into a single noise vector by summing up their counts in each droplet (**Fig. 3D**). GUMM then performs principal component analysis (PCA) on the allele frequencies of the common alleles and the noise vector. A multivariate GMM is then fit to the first two principal components (PCs) of the data to identify droplet clusters. Hartigan’s dip test was performed on the first PC to determine K. If K=2, ***μ***_*k*_ is a vector containing the PC means and **Σ**_*k*_ is the covariance matrix of the PCs. The determinant of Σ is computed for each cluster to identify the one with highest droplet density (smallest determinant). The cluster with higher density in low dimensional space corresponds to true heterozygous droplets:

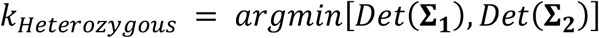

The skewed univariate, univariate, and multivariate GMMs were implemented using the *mixsmsn* and *mclust* R packages, respectively^12,13^.

### Analysis of public scDNAseq data

scDNAseq data from ten Hacken et al. was downloaded from Sequence Read Archive (SRA) under Bioproject accession number PRJNA665752. The data included a Ba/F3 sample consisting of an admixture of cells edited at only one of the following loci: *Trp3, Birc3, Atm, Chd2, Mga*, or *Samhd1*. It also included a multiplexed Ba/F3 sample where multiple edits were present in the same cell. Both samples were processed using the same procedure and parameters as with the artificial cocktails except reads were instead aligned to Gencode’s mm10 (GRCm38.p4) mouse reference genome and CRISPResso2 was run using WGS mode with a flank parameter of 15bp. In the singleplexed sample, droplets with less than 120 total reads, corresponding to approximately 20 reads per gene, were removed from the CRISPResso2 output.

### Data Access

Processed data and code are deposited in Zenodo under DOI:10.5281/zenodo.15398817.

## Results

### Generation of a gene editing scDNAseq data resource

To create our data resource, we employed CRISPR-Cas9 to modify HL-60 cells at *CLEC12A*. Single-plex and multi-plex edited cells were isolated and expanded to create HL-60 monoclonal cell lines (**Figure 1A**). Sanger sequencing and ICE analysis^14^ identified several single-plex edited clones edited at *CLEC12A* including one that was homozygous with a +1 insertion (+1Ins) and one that was heterozygous with a -8del and -9del compound deletion (**Figure 1B**).

**Fig. 1:**
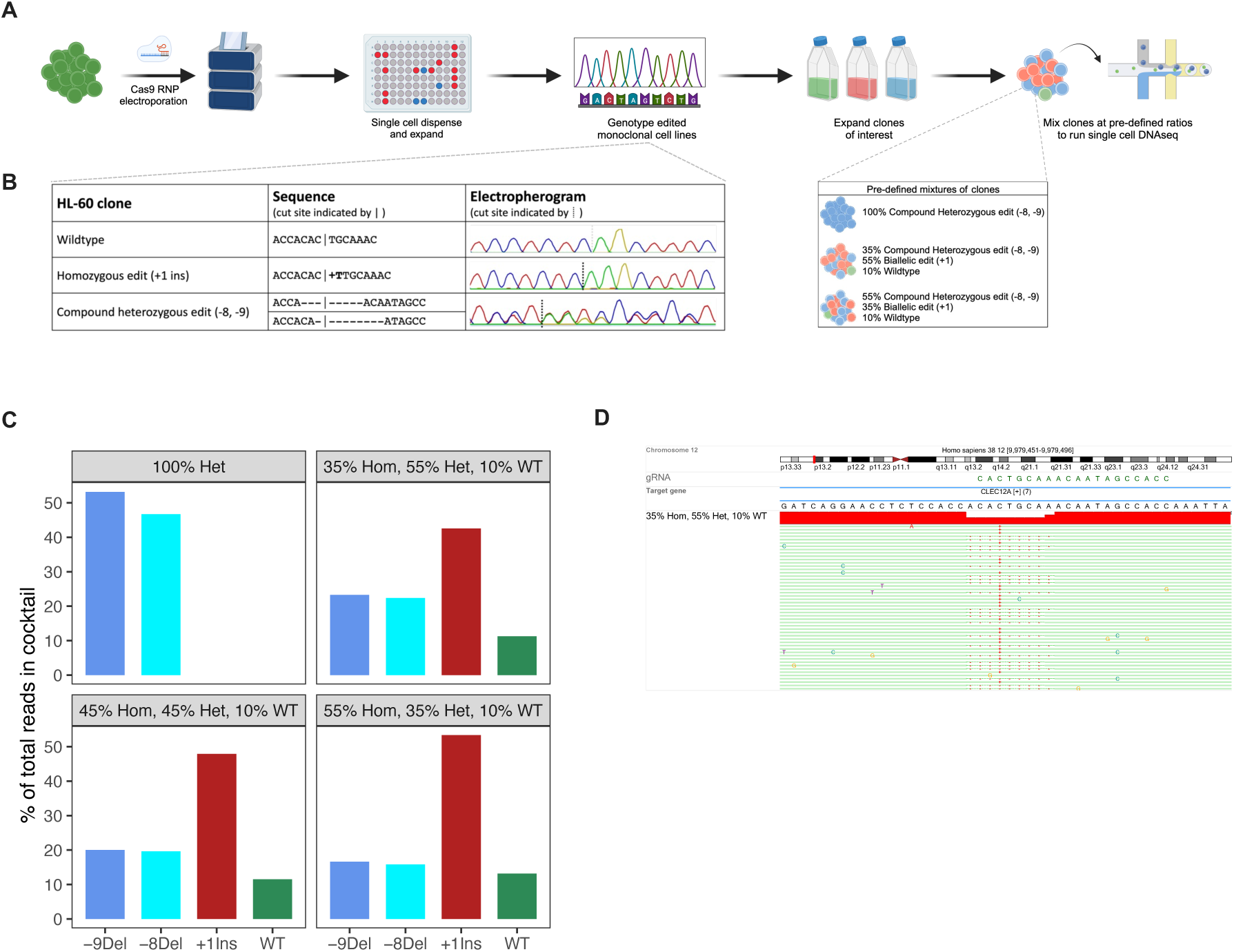
Generation and validation of artificial cocktails (**A**) Graphical overview of procedure used to generate artificial mixtures of edited HL60 monoclones (**B**) Electropherogram from Sanger sequencing of WT HL60 and two monoclones with homozygous and heterozygous editing of *CLEC12A*. Vertical line indicates cut site of CRISPR-Cas9. (**C**) Electropherogram from Sanger sequencing of HL60 clone harboring edit at *CLEC12A*. Vertical lines indicate cut sites. (**D**) Bar plots showing total number of reads mapping to the four possible alleles across three cocktails.

The single-plexed clones were mixed at varying ratios to generate three artificial cocktails. The first comprised a mix of homozygous, heterozygous, and WT clones in proportions of 35%, 55%, and 10%, respectively. This sample was prepared with an additional filtering step that minimized the multiplet rate (See methods). Two more cocktails were created using the same clones but in proportions of 55%, 35%, and 10% and 45%, 45%, and 10%. Prior to library generation, the first cocktail was subjected to an additional filtering step whereas the last two were not in order artificially produce both clean and noisy data with varying multiplet rates. We then employed Mission Bio’s Tapestri platform to generate barcoded single-cell DNA libraries for these two cocktails and the single-plexed compound heterozygous clone (**Figure 1A**). After sequencing these libraries, we analyzed the allele composition at the pseudobulk level and observed allele frequencies consistent with the makeup of the cocktail (**Figure 1D**).

### Exploring artifacts that confound allelism interpretation

Because the single-plexed cocktails are comprised of pure clones with pre-defined editing patterns at the *CLEC12A* locus, the scDNAseq readout of allele frequencies should theoretically reveal only droplets with 100% WT reads, 100% of reads containing the +1Ins allele, or reads equally distributed between the -8Del and -9Del patterns. Indeed, we observed these genotypes in our data but noticed several other interesting artifacts (**Figure 2A**). First, we observed droplets with more than two unique alleles indicating that two cells were encapsulated within the same droplet. Since identifying droplets with ploidy greater than two, assuming a human context, is relatively straightforward we refer to them as transparent multiplets (**Figures 2B and 2C**). We also identified droplets that displayed an allele frequency distribution consistent with that of a heterozygous cell but whose genotype cannot exist due to the clonal nature of our cocktails (e.g. WT/+1Ins) (**Figures 2B and 2C**). Because identification of these multiplets is less straightforward we refer to them as opaque multiplets.

**Fig. 2:**
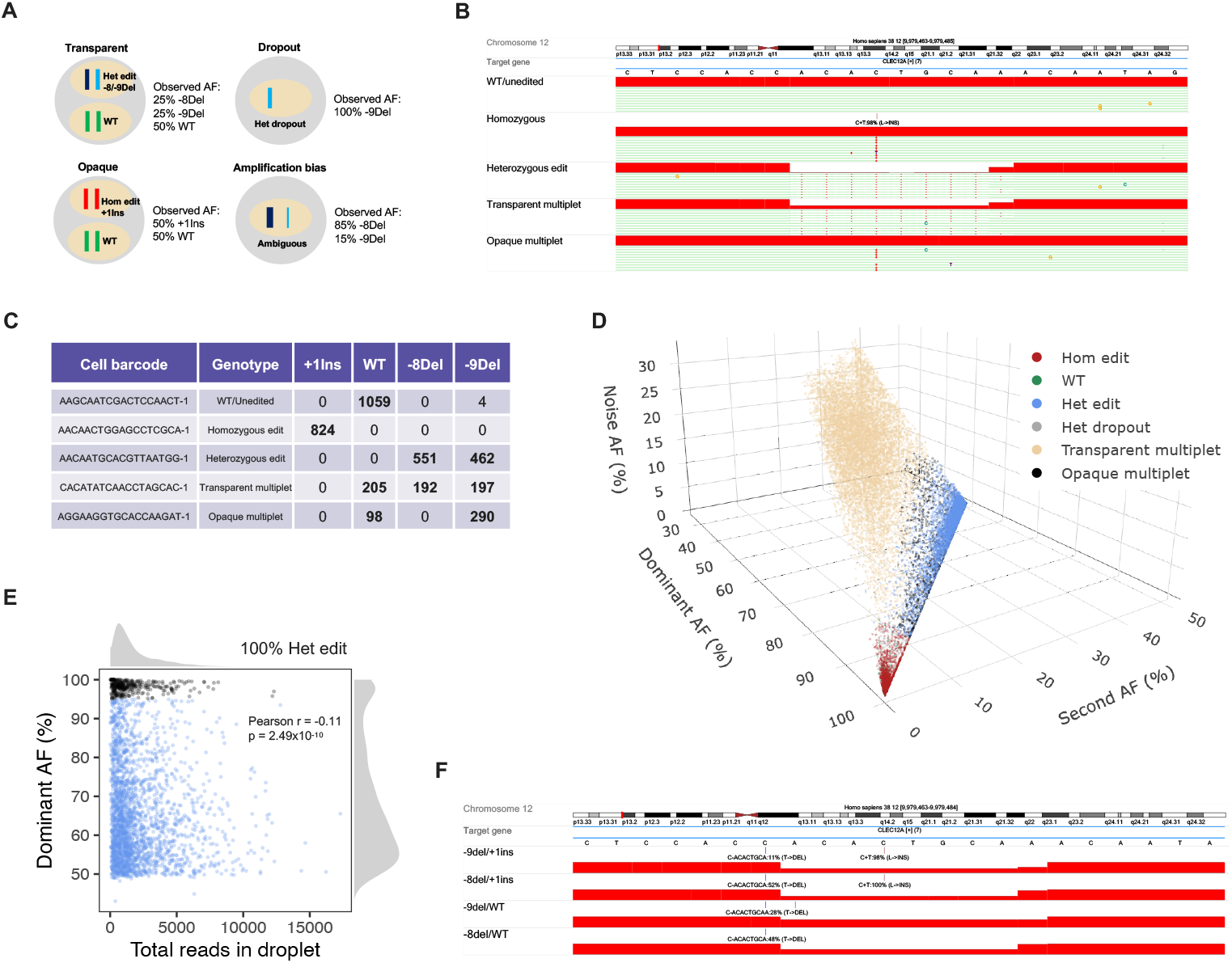
Single-cell characterization of artificial cocktails (**A**) Illustration of possible artifiacts in scDNAseq data and their resulting allele frequency readouts that confound interpretation of gene editing allelism. (**B**) Read alignment profiles of six selected droplets from cocktails displaying different allele combinations observed at the *CLEC12A* edit site. (**C**) Table of allele read counts produced by CRISPResso2 for the six droplets. (**D**) 3D scatterplot showing frequencies of each droplet’s top three most frequent *CLEC12A* alleles across all cocktails. (**E**) Scatterplot showing relationship between read depth and dominant allele frequency of droplets from sample with 100% compound heterozygous edited cells. Read depth is the total number of reads in the droplet and dominant allele is the allele with the most reads mapped to it in the droplet. Black points indicate droplets with a dominant allele frequency of >95% which are considered dropouts (**F**) Read alignment profiles of four opaque multiplets with highly unlikely allele combinations.

Second, our data show that heterozygous allele counts typically deviate from a theoretical 50-50 frequency distribution, often displaying a bias towards one allele (**Figure 2D, Figure 2E**). This is most likely attributed to amplification bias, which arises during PCR amplification of minimal starting DNA quantities^8^, as is the case with droplet-based amplification. While this phenomenon is anticipated in microfluidic technologies, it introduces uncertainty when determining heterozygosity of an edited cell. In addition, we found that droplets with higher read depth at the *CLEC12A* editing site ameliorated some of this bias and improved the separability of homozygous and heterozygous droplets when analyzing their allele frequencies (**Figure S1C**). Minor pre-processing of droplet allele frequencies also improves the ability to resolve genotypes. For example, the dominant allele frequency of homozygous cells can approach 100% if we remove alleles with less than 10 reads (**Figure S1**). This filtering is possible with our data since we deeply sequenced our libraries to 250 reads per droplet per amplicon on average (**Table S1**). Nonetheless, this solution may not be viable for all scDNAseq experiments, as shown by prior studies that achieved an average sequencing depth of around 20 reads per droplet per amplicon^6^. Thus, setting stringent filtering criteria may be counterproductive when analyzing less deeply sequenced libraries.

Moreover, we observed allele dropout in our heterozygous clone sample where either the -8Del or -9Del allele was nearly undetectable from the readout suggesting that this cell would have been categorized erroneously as homozygous. This occurred in 12.3% of the droplets and is within the expected range for this platform. There was also a weak but significant association between dropout rate and droplet read depth (Pearson r = -0.11, p = 2.49×10^−10^) (**Figure 2E, Figure S1C**). Multiplets and dropouts were not mutually exclusive as we observed droplets with genotypes including +1Ins/-8Del, +1Ins/-9Del, WT/-8Del, and WT/-9Del. These erroneous genotypes are consistent with a scenario where two cells are encapsulated in the same droplet with one cell experiencing allele dropout or amplification bias (**Figure 2F**).

### Automated genotyping with Gaussian mixture models

Since these artifacts are difficult to resolve and may occur at varying frequencies between experiments, previous studies have chosen a simplified approach to genotyping cells by setting hard allele frequency thresholds to determine heterozygosity^6^. These strict cutoff criteria can vary both across different experiments and among various pre-processing steps which can hinder consistent and reproducible analysis. Furthermore, no study has proposed a standardized procedure for identifying and handling multiplets from scDNAseq of gene edited cells. Existing methods have focused on identifying multiplets from an admixture of cells with distinct mutational profiles across several different loci^11^.

However, many gene editing experiments involve editing a pure population of cells all at the same loci making edited allele variants the only differentiator between individual cells. To address these limitations, we developed a systematic computational workflow named GUMM that performs automated artifact-aware genotyping by fitting a series of Gaussian mixture models to the allele frequency readout of single droplets in a stepwise fashion. GUMM involves three major steps which include identifying homozygous droplets, flagging transparent multiplets, and distinguishing heterozygous droplets from opaque multiplets. (**Figure 3A**).

**Fig. 3:**
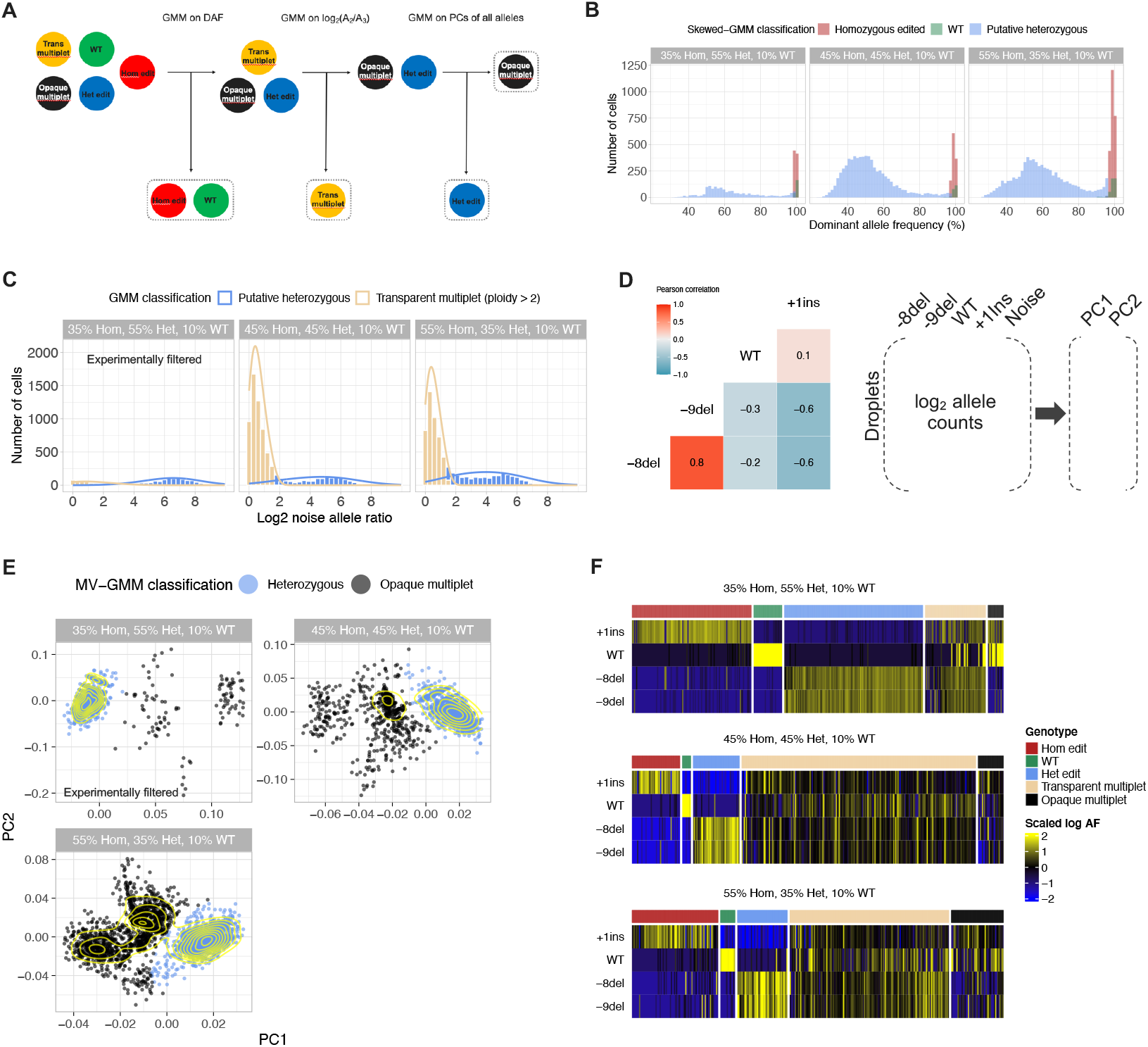
Automation of single-cell genotyping using GUMM (**B**) GUMM workflow which automates both genotyping and multiplet detection by fitting three separate Gaussian mixture models in series. (**B**) Histograms showing frequency of the dominant *CLEC12A* allele in droplets for each genotype class predicted by fitting a skewed univariate GMM (**C**) Histograms of log2 noise allele ratios for droplets with >2 detected *CLEC12A* alleles. Color indicates droplets predicted to be transparent multiplets or putative heterozygous edited droplets after fitting a univariate GMM. (**D**) Heatmap showing correlation of each of four possible *CLEC12A* alleles across putative heterozygous droplets. PCA was used to reduce log2 read frequencies of all alleles into two dimensions. Read frequencies of rare alleles are summed into a single “noise” feature prior to dimensionality reduction. (**E**) Scatterplots with density contours showing putative heterozygous cells in low dimensional space with distinct clusters corresponding to true heterozygous singlets and opaque multiplets. Color shows automated classification of clusters by fitting a multivariate GMM to PCs. (**F**) Heatmaps showing scaled log frequencies of the four *CLEC12A* alleles across droplets. Top annotation bar indicates the *CLEC12A* genotype category each droplet was assigned to by GUMM.

To evaluate the performance of GUMM, we applied it to our ground truth scDNAseq data generated from our artificial clonal cocktails starting with samples containing singplexed edits. In the first step, GUMM successfully classified droplets into two distinct populations corresponding to homozygous and putative heterozygous droplets by fitting a skewed univariate GMM to read counts of the dominant *CLEC12A* allele in each droplet (**Figure 3B**). No pre-filtering of low count alleles was performed in droplets to demonstrate that GUMM is robust to amplification bias and sequencing error. The same analysis was performed on pre-filtered data with similar results (**Figure S1B**). Moreover, GUMM calculates the probability of each droplet’s membership within the identified clusters, enabling the exclusion of droplets with genotype predictions of low confidence (**Figure S2**). In the second step, GUMM leverages the ploidy information of each droplet to identify those that clearly have more than two *CLEC12A* alleles. The ploidy of a droplet is first summarized by taking the log_2_ ratio between the counts of the second most common *CLEC12A* allele and those of the third most frequently occurring *CLEC12A* allele, also referred to as the noise allele. The distribution of this noise allele ratio statistic unveils two distinct populations both of which were effectively clustered by GUMM (**Figure 3C**). The cluster with the lowest average log_2_ noise allele ratio corresponds to transparent multiplets and display a triploid allele frequency distribution (**Figure 3C**). The remaining pool of unclassified droplets are comprised of true heterozygous cells and opaque multiplets which occur when two homozygous cells (e.g. WT/+1ins) are enclosed in the same droplet yielding an allele frequency distribution consistent with that of a heterozygous cell. In order to distinguish these multiplets from true heterozygous cells, we postulated that true heterozygous cells would consist of alleles that co-occur frequently across droplets. Indeed, we observed this association in our ground truth data where the -8Del allele is strongly correlated with the -9Del allele (Pearson r = 0.81, p << 0.01) (**Figure 3D**). Likewise, the heterozygous alleles are anti-correlated with the +1ins and WT alleles (**Figure 3D**). GUMM captures this correlation structure in the remaining droplet pool by implementing principal component analysis (PCA) on the allele counts of the most frequently occurring alleles and the collapsed noise feature (See methods for details). On a dataset where homozygous and heterozygous alleles are not known *a priori*, we sum the counts of all alleles and set a loose frequency cutoff (e.g 1-5%) to select common alleles for PCA. As the final step, GUMM fits a multivariate GMM to the first two principal components (PCs) to effectively cluster heterozygous cells separately from opaque multiplets (**Figure 3E**). Inspection of each droplet’s predicted genotype and their corresponding allele frequencies shows consistency with ground truth (**Figure 3F**). In contrast, simple hierarchical clustering of the allele frequencies cannot achieve the same accuracy and resolution of GUMM (**Figure S3**).

After determining allelism at the single-cell level and identifying multiplets, GUMM estimates the sample composition by tallying the droplet genotypes after removing multiplets or splitting them into partial droplets (See methods for details). In our analysis, we removed all multiplets and found that GUMM was able to accurately estimate the original clonal fractions of the cocktails with less than 10% deviation in the 35% Hom, 55% Het, 10% WT cocktail and less than 5% deviation in the 55% Hom, 35% Het, 10% WT and 45% Hom, 45% Het, 10% WT cocktails (**Figure 4B**). Although the 35% Hom, 55% Het, 10% WT cocktail contained less multiplets, we observed a slightly larger deviation from ground truth. Closer inspection of this sample revealed xxx heterozygous dropouts that were classified as homozygous. While our genotype composition estimate closely approximates the ground truth, we expect that the homozygous fraction of a sample will consistently be overestimated, and the degree of this overestimation will likely be related to heterogeneity and allele dropout rate.

**Fig. 4:**
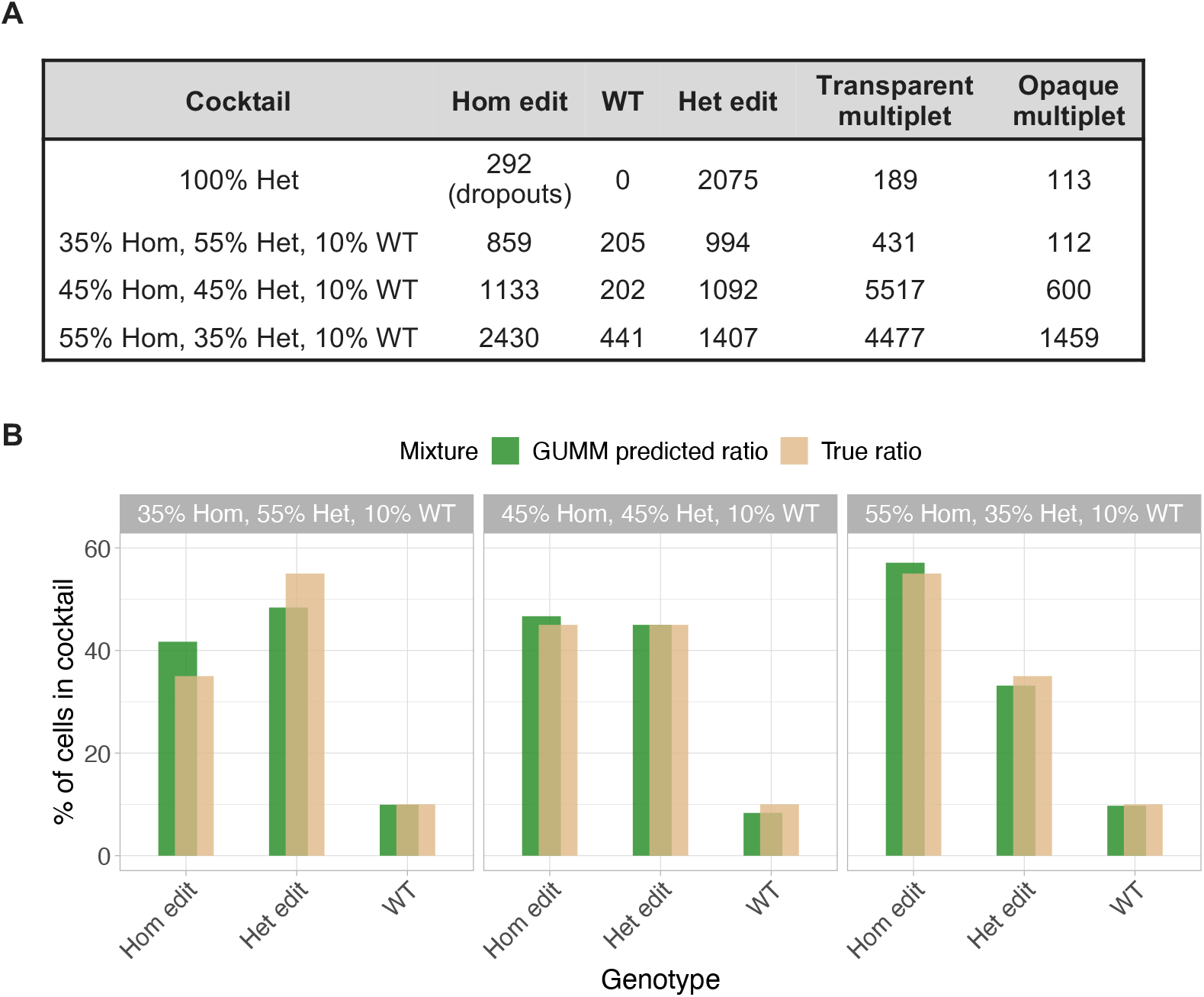
Estimating clonal composition of artificial cocktails (**A**) Table of droplet counts in each genotype category across cocktails. (**B**) Bar plots comparing sample composition estimated by GUMM with ground truth for the 35% Hom, 55% Het, 10% WT, 45% Hom, 45% Het, 10% WT, and 55% Hom, 35% Het, 10% WT cocktails.

### Analysis of public scDNAseq gene editing data

To demonstrate that GUMM can be applied across different datasets we applied it to a published scDNAseq dataset generated from Ba/F3 mouse cells edited across six genes (*Atm, Birc3, Chd2, Mga, Samhd1, Trp53*) with CRISPR-cas9. The first sample consisted of an admixture of singleplex-edited Ba/F3 cells which enabled an orthogonal approach for identifying multiplets. Namely, cells harboring multiple edited loci are likely to be multiplets. In this sample, GUMM was able to identify transparent multiplets but not opaque multiplets due to the allelic heterogeneity and the relatively low composition of diploid droplets, violating the assumption that most heterozygous cells comprise of two co-occurring alleles. Despite this limitation, we were still able to predict sample genotype composition across the six genes and found that our estimates were concordant with the published results which adopted hard allele frequency threshold cutoffs (**Figure 5A**). Since this sample consists of primarily WT cells at any given gene, we compared the WT cell composition estimates with the published results and observed strong correlation across the six genes (Spearman r = 0.94, p = 0.02) (**Figure 5B**). We compared the two orthogonal approaches for multiplet identification and found that GUMM flagged 46.4% of cells with multigene edits as transparent multiplets and 95.1% of cells with a single gene edit as singleplets. Moreover, we increased the stringency of a droplet being identified as transparent multiplet by requiring evidence from at least two genes. At this stringency we found that 19.7% of droplets with multigene edits were flagged as transparent multiplets and nearly 100% of droplets with editing at one gene were labeled as singleplets. These results suggest that in a true gene editing experiment where all cells are intended to harbor edits at the same genes, GUMM will still be able to identify a significant portion of multiplets in the data by just analyzing allele variants at each individual locus.

**Fig. 5:**
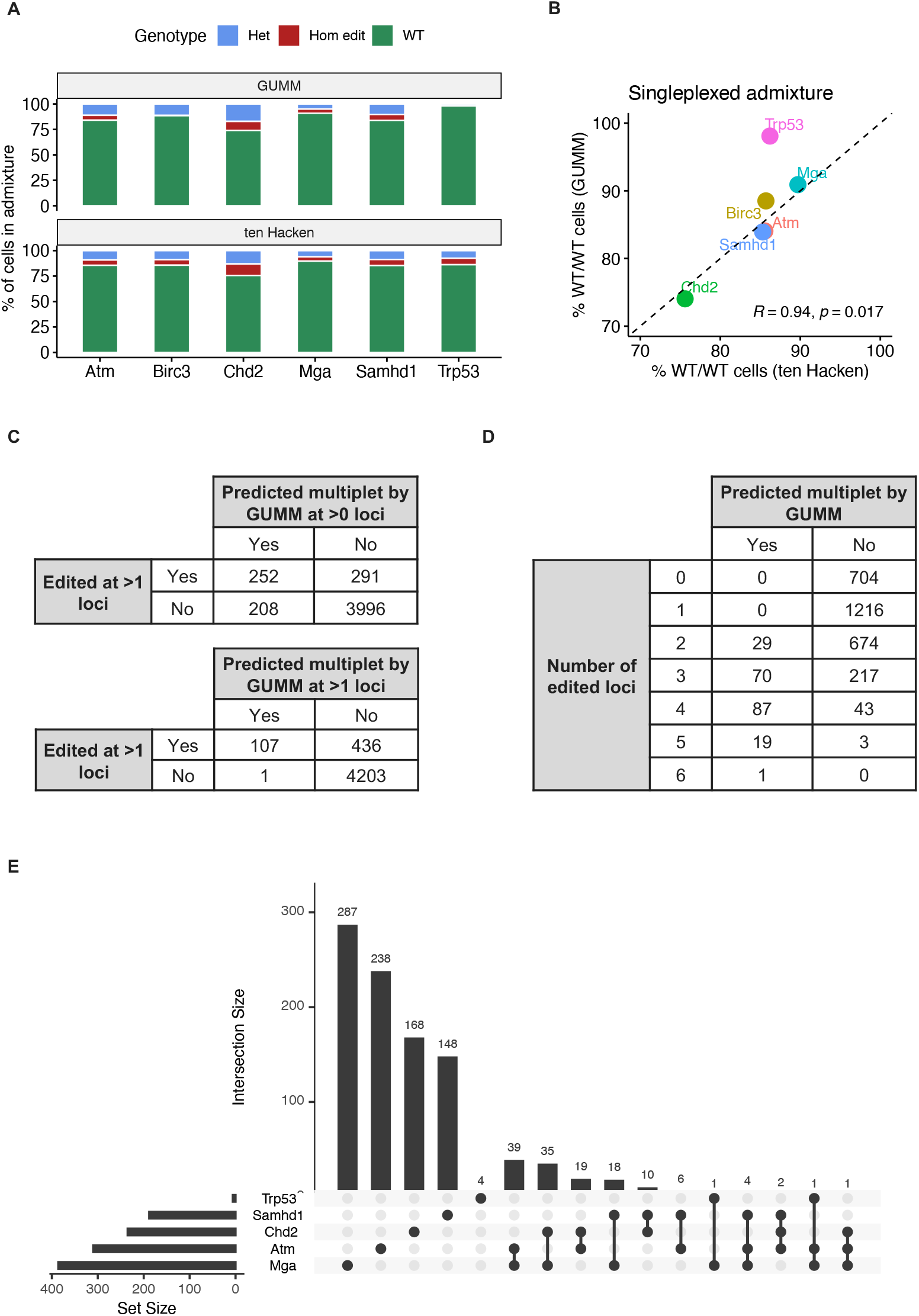
Application of GUMM to public scDNAseq data (**A**) Comparison of reported sample composition with GUMM’s estimate from scDNAseq of singleplexed Ba/F3 sample published by ten Hacken et al. (**B**) Scatterplot showing correlation between the percent of WT/WT (unedited) cells estimated by GUMM and the percent reported in original study. Black diagonal dotted line indicates perfect concordance (**C**) Tables showing overlap between droplets being flagged as transparent multiplets by GUMM and those with editing at more than 1 gene in the singleplexed Ba/F3 sample. (**D**) Table showing the overlap between droplets identified as multiplets by GUMM and those with simultaneous edits across the six genes in the multiplexed Ba/F3 sample. (**E**) Upset plot showing number of cells with different combinations of homozygous gene edits.

Next, we applied GUMM to the multiplexed sample data where Ba/F3 cells were transduced with a pool of lentivirus expressing sgRNAs to simultaneously edit the six genes. As expected, we found that droplets with no editing or editing at just one gene were unlikely to be flagged as a potential multiplet by GUMM (**Figure 5D**). Droplets harboring several edited genes were more likely to be flagged as a multiplet. Since it is less likely for any given cell to absorb more than one sgRNA, these results suggest that these droplets most likely contain multiple singleplex edited cells rather than encapsulating a single true multiplex-edited cell (**Figure 5D**). After removing multiplets, we found that only 136 out of 3063 cells contained homozygous edits at more than one gene suggesting that a pooled lentivirus approach may not be ideal for efficient multiplex editing (**Figure 5E**).

## Discussion

In this study, we performed a deep and comprehensive analysis of “ground truth” scDNAseq of artificial gene edited cocktails. We leveraged these cocktails to explore artifacts that could misguide interpretation of allelism when applying this technology to gene editing. These include amplification bias of alleles and multiplets which we show can be identified and accounted for by applying GMMs to transformations of the allele frequency readout. Because most gene editing experiments in the context of cell therapy are intended to introduce identical edits to each cell, most cells will be edited at the same loci making multiplet identification only possible through deep analysis of allele variants. With these concepts in mind, we developed a standardized workflow called GUMM to implement our approach in an automated and unbiased fashion. We show that our approach can accurately genotype cells and estimate the original ratios of our cocktails regardless of the multiplet rate.

We found that when applied to previously published data, GUMM was unable to identify opaque multiplets but was still able to flag a substantial fraction of transparent multiplets by analyzing only the allele variants. We believe this was due to the allelic heterogeneity of the putative heterozygous droplets caused either by inconsistent editing or additional noise we could not remove. This suggests that GUMM is best suited for analyzing scDNAseq from gene editing experiments employing gRNAs that generate a constrained set of indels. Consequently, we anticipate that GUMM may encounter challenges in detecting opaque multiplets in scDNAseq data obtained from cells edited with base editing, given the extensive pool of potential allele variants stemming from bystander modifications. Nevertheless, with the emergence of prime editing as an alternative technology characterized by fewer bystander modifications^15^, we envision an enhanced utility for GUMM. Our study provides both a rich data resource and computational workflow for analyzing scDNAseq data from gene editing experiments.

