## Supplementary figures and images for "A resource and computational approach for quantifying gene editing allelism at single-cell resolution"

### Figure S1

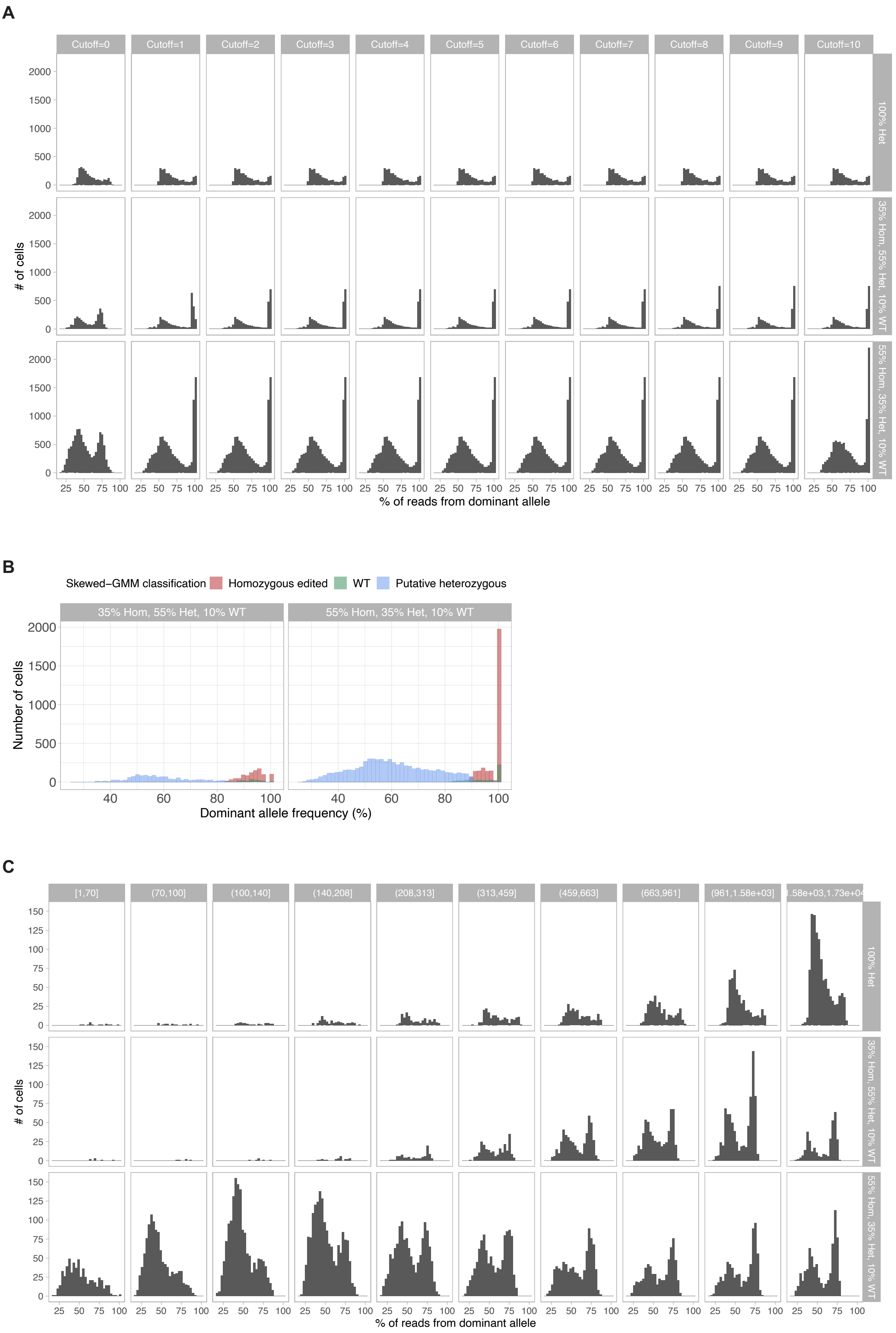

### Figure S2

**A**

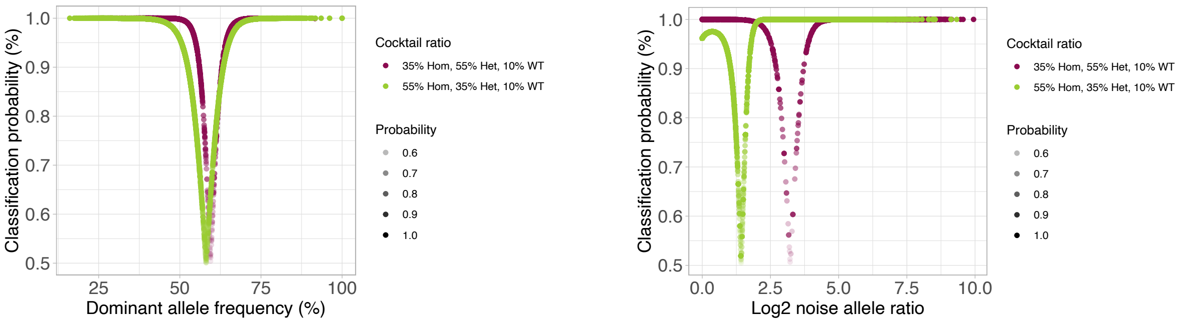


**B**

| | | Predicted multiplet by GUMM at >1 loci | |
|---|---|---|---|
| | | Yes | No |
| Edited at >2 loci | Yes | 107 | 436 |
| | No | 1 | 4203 |

### Figure S3

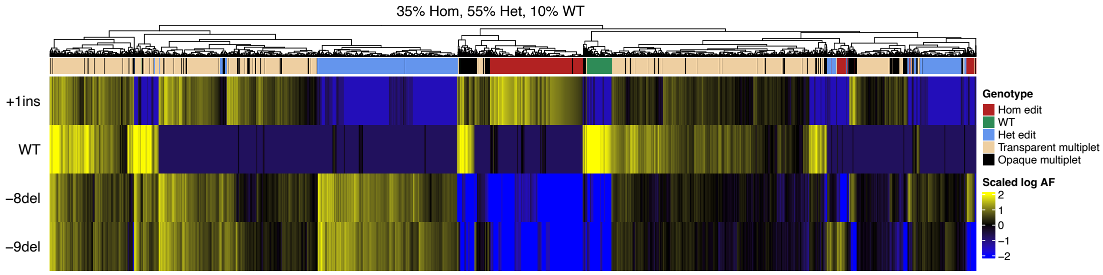

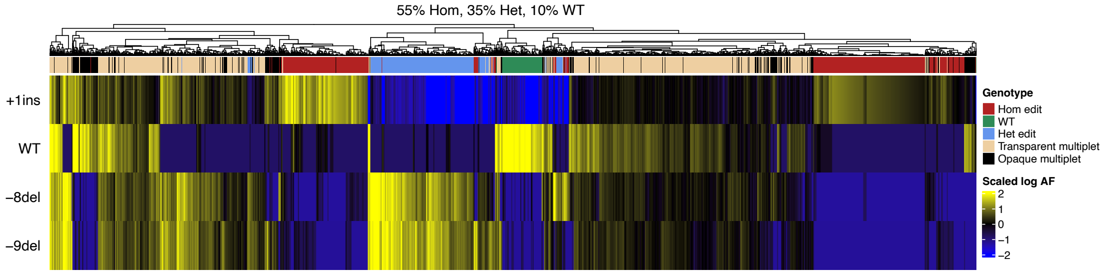
